# Two frequency bands contain the most stimulus-related information in visual cortex

**DOI:** 10.1101/049718

**Authors:** Christopher M. Lewis, Conrado A. Bosman, Nicolas M. Brunet, Bruss Lima, Mark J. Roberts, Thilo Womelsdorf, Peter de Weerd, Sergio Neuenschwander, Wolf Singer, Pascal Fries

## Abstract

Sensory cortices represent the world through the activity of diversely tuned cells. How the activity of single cells is coordinated within populations and across sensory hierarchies is largely unknown. Cortical oscillations may coordinate local and distributed neuronal groups. Using datasets from intracortical multi-electrode recordings and from large-scale electrocorticography (ECoG) grids, we investigated how visual features could be extracted from the local field potential (LFP) and how this compared with the information available from multi-unit activity (MUA). MUA recorded from macaque V1 contained comparable amounts of information as simultaneously recorded LFP power in two frequency bands, one in the alpha-beta band and the other in the gamma band. ECoG-LFP contained information in the same bands as microelectrode-LFP, even when identifying natural scenes. The fact that information was contained in the same bands in both intracortical and ECoG recordings suggests that oscillatory activity could play similar roles at both spatial scales.

## Introduction

Visual stimulus representation depends on recurrent and lateral connectivity in populations of visual cortical neurons. Recurrent connectivity, especially of local inhibitory cells, gives rise to oscillatory synchronization. The oscillatory synchronization of pairs of visual cortical neurons has been found to depend on the specific characteristics of visual stimulation, such as co-linearity, common fate and other Gestalt principles. Likewise, rhythmic neuronal synchronization in specific frequency ranges has been found to reflect stimulus properties like contrast, velocity, size and eccentricity (Engel et al., 1990; Gray et al., 1989; Singer and Gray, 1995), as well as cognitive variables, such as attention (Fries et al., 2001). Understanding how visual features, at one end, and whole visual scenes, at the other, are represented by the population activity of visual areas is essential for improving our understanding of sensory coding. Such knowledge can help us develop realistic schemes for sensory processing, as well as enhance our ability to read-out brain activity in real time, in order to develop brain-machine interfaces (BMIs).

Local oscillatory activity can also be measured in the aggregate synchronization of sub-and supra-threshold neuronal events, as captured by the power of band-limited components in the LFP. Specific frequency bands of invasively and non-invasively recorded electrical activity have likewise been shown to reflect aspects of visual stimulation, such as categorical distinctions (Gonzalez Andino et al., 2007), or low-level stimulus features (Belitski et al., 2008; Berens et al., 2008; Besserve et al., 2015; Jia et al., 2011; Kayser and König, 2004; Myers et al., 2015). Likewise, oscillatory power recorded from visual (Engel et al., 1990; Gray et al., 1989; Rotermund et al., 2013; 2009; Singer and Gray, 1995) or frontal cortex (Fries et al., 2001; Tremblay et al., 2015), can indicate the locus of visual attention on a trial-by-trial basis. These findings suggest that the LFP may reflect local processing of sensory signals, as well as provide a robust and reliable signal for BMIs.

We sought here to systematically assess the degree to which oscillatory synchrony in the LFP reflects characteristics of visual stimulation across a broad range of stimulus properties: orientation, direction of motion, absolute stimulus position, eccentricity and polar angle, as well as the identity of natural images. In intracortical microelectrode recordings, we directly compared the amount of information related to low-level visual features (orientation and direction of motion) retrievable from LFP power and simultaneously recorded spike-rate estimates from V1 populations. We found that comparable amounts of information were present in the spike rate and the local pattern of oscillatory power in two frequency bands: the alpha-beta band and the gamma band. In subdural electrocorticography (ECoG) recordings, it was possible to decode stimulus position as well as the identity of natural images in the same frequency bands, suggesting that the pattern of oscillatory synchrony across early visual areas may reflect global aspects of stimulation. By combining data from different recording techniques and viewing conditions, we were able to demonstrate that the same two frequency bands robustly indicate both low-level and holistic features of visual stimulation.

## RESULTS

We investigated the extent to which characteristics of visual stimulation were recoverable in specific frequency bands of the LFP. In order to evaluate the generality of our findings, we combined intracortical multi-electrode recordings from primary visual cortex of 3 awake, behaving monkeys with recordings from large-scale ECoG grids covering large portions of one hemisphere in 3 additional, behaving monkeys. Across a variety of stimulus features, viewing contexts, recording techniques, and 6 monkeys, we found that power in two frequency bands reliably had the most stimulus-related information. These two bands, centered between 10-20 Hz and 40-120 Hz exhibited distinct temporal dynamics that were reproducible across recording techniques, decoded feature and stimuli. The 10-20 Hz band exhibited fast, transient performance that was largely attenuated during the sustained period of stimulation. In contrast, power in the band between 40-120 Hz showed both fast, transient performance, as well as sustained performance through the duration of the stimulus.

### Time-frequency dynamics of orientation and direction information and classification error

In order to evaluate the quality of stimulus-related information carried by the LFP, we investigated the representation of orientation and direction from localized moving gratings. To do this, we first computed single trial estimates of spectral power in a time-resolved manner. We then applied naïve-Bayes classification to our single-trial power estimates in order to produce a prediction of the desired feature given the power at a specific point in time and for a specific frequency. This resulted in training a single decoder for each point in the time-frequency plane, which could estimate stimulus position based on the spatial pattern of LFP power at a given frequency. The details of the decoder we used and how we applied it are available in the Materials and Methods section. We were able to compare the information available in the LFP to the spiking activity of small groups of neurons recorded from the same electrodes. We found that two frequency bands carried the most information related to both the orientation and direction of moving gratings. The temporal dynamics of the two bands were distinct, with the alpha-beta band showing a phasic increase in information immediately following stimulation and the gamma band showing sustained information throughout stimulus presentation (Fig. 1A shows information about stimulus orientation, averaged across 53 sessions in 3 monkeys (Monkey J: 21 sessions, Monkey L: 26 sessions, Monkey N: 6 sessions), chance = 1/8 (12.5%); Fig. 1C shows information from the same data about the direction of stimulus motion, chance = 1/16 (6.25%)). While some individual sessions had comparable amounts of information in the LFP and the MUA (Fig. 1C, F), across all sessions, the accuracy of the decoder based on gamma band power was about 5% less in absolute terms than that based on MUA. Fig. 1B and D compare orientation and direction decoding based on MUA, LFP gamma power, alpha-beta power and the broadband LFP amplitude for the same sessions as Fig. 1A and C. From each session, we used the same number of MUA and LFP channels, so that the number of features was identical across decoders. In addition, we limited analysis to sessions with at least 3 electrodes containing both MUA and LFP (range = 3-9, mean = 4.2 electrodes).

**Figure 1.**
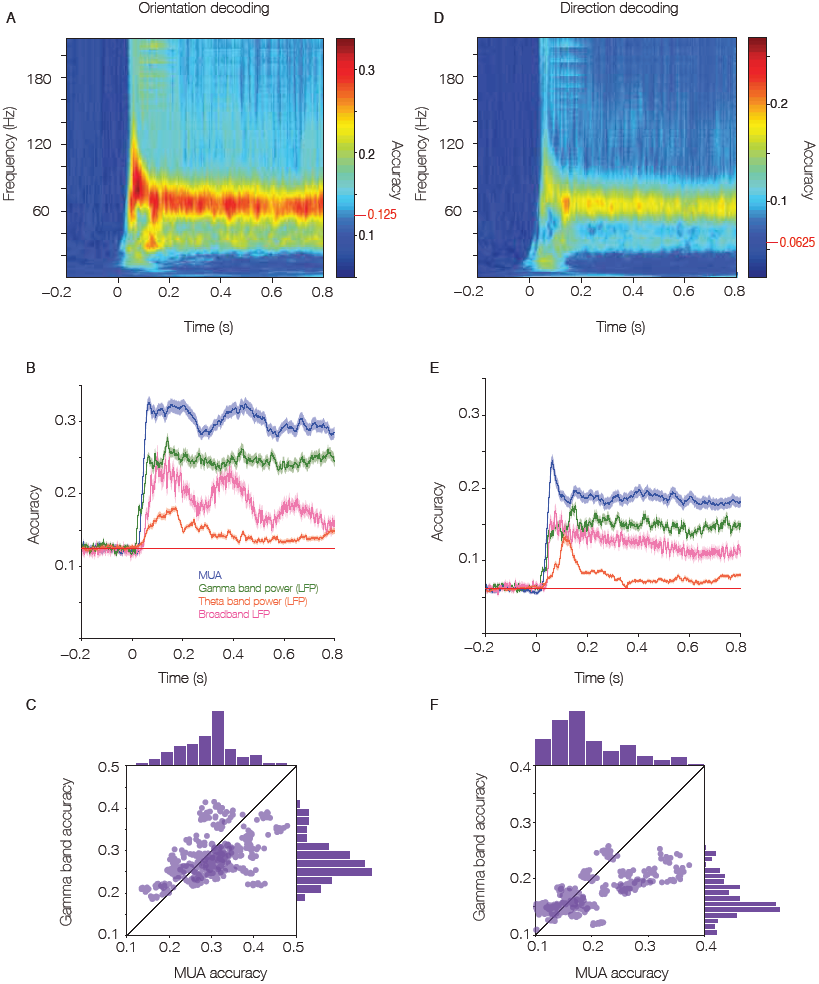
Orientation and direction decoding from LFP and MUA are comparable. Comparison of average LFP and MUA decoding performance across sessions and animals. (A) Time-frequency classification performance for a decoder trained separately on each time-frequency point to distinguish orientation based on intracortically recorded LFP. The red line indicates chance level performance. (B) Direct comparison of the accuracy for decoders trained on the same channels as shown in A, but using the multi-unit response (blue), the gamma band (60-90 Hz) power (green), the alpha-beta power (orange), or the broadband voltage (pink). The shaded region depicts the 95% confidence interval across sessions. The red line indicates chance level performance. (C) Comparison of performance of decoding orientation based on Gamma band power and MUA rate estimates. (D) Same as A, but for direction of motion. (E) Same as B, but for direction of motion. The shaded region depicts the 95% confidence interval across sessions. All results are average performance across multiple sessions in 3 monkeys. (F) Same as C, but for direction of motion.

Given that the LFP and MUA activity contained stimulus information in such a specific manner, we next investigated the size of errors made during misclassifications. The distributions of error sizes across the trials of one example session for both LFP and MUA decoding of orientation are displayed in Fig. 2A and B. Both the LFP and MUA are able to classify the orientation of a given stimulus with high accuracy, and errors are in general clustered around the veridical orientation. The distribution of error sizes across the trials of the same session for both LFP and MUA decoding of direction are displayed in Fig. 2C and D. Like for orientation, errors in the estimation of direction from LFP and MUA are both small, with the interesting fact that LFP shows a higher incidence for decoding the movement in the opposite direction. Finally, we quantified the error in orientation decoding across the population of sessions and monkeys in a time-resolved fashion, as the average distance in radians from the veridical stimulus orientation. The time resolved average error for LFP is shown in Fig. 2E and that for MUA is shown in Fig. 2F. Errors were in general less than 1.75 radians for LFP and 1.5 radians for the MUA.

**Figure 2.**
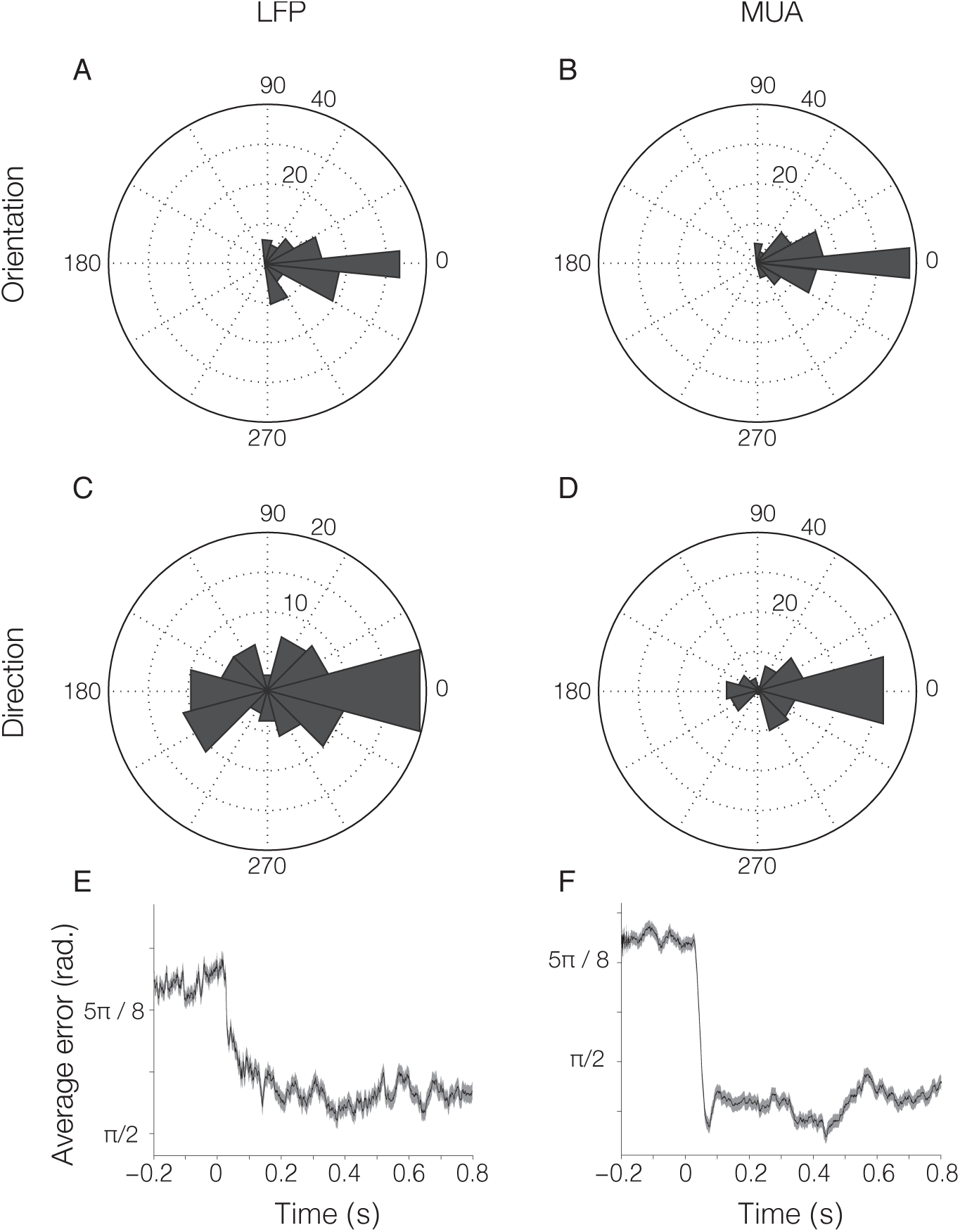
Misclassification errors are small and comparable between LFP and MUA. (A) Polar plot showing the proportion of classified orientations (true stimulus orientation aligned to 0) using LFP in the gamma band. (B) Same as in A, but using spike density. (C) Same as A, but for stimulus direction. (D) Same as B, but for stimulus direction. Errors made in classifying the orientation of stimuli from the LFP and MUA were highly similar, as shown in Fig. 2A and B. In the case of direction, the LFP decoder exhibited a higher tendency to misclassify stimuli as occurring in the opposite direction as compared to MUA. (E) Mean error distance for LFP decoding of orientation as a function of time. The average distance between the veridical stimulus orientation and the estimated orientation is shown in radians. Shaded regions show 95 percent confidence interval across sessions and monkeys. (F) As (E), but for MUA decoding.

### Comparison of LFP versus MUA on a single-trial basis

Next, we assessed the degree to which the decoded stimulus orientation co-varied between MUA and LFP on a trial-by-trial basis, and as a function of time after stimulus onset. The circular correlation across single trials, between stimulus orientation decoded from LFP and MUA is shown in Fig. 3A as a time-frequency plot around stimulus onset. The observed correlation is specific to the frequency bands showing the highest degree of stimulus specificity, though the absolute value of correspondence is moderate, suggesting that independent information may be carried by the two measures of local activity. Despite the moderate correlation in decoded stimulus orientation, errors in MUA and LFP decoding co-occurred relatively frequently, perhaps due to a common dependence on the animal’s arousal. Fig. 3B displays the proportion of joint error trials as a function of time. The green curve shows that trials that resulted in an error when decoding orientation from the LFP also resulted in an error when decoding based on the MUA 60-70% of the time. The blue curve shows that trials that resulted in an error when decoding the orientation from the MUA also resulted in an error when decoding based on the LFP 75% of the time.

**Figure 3.**
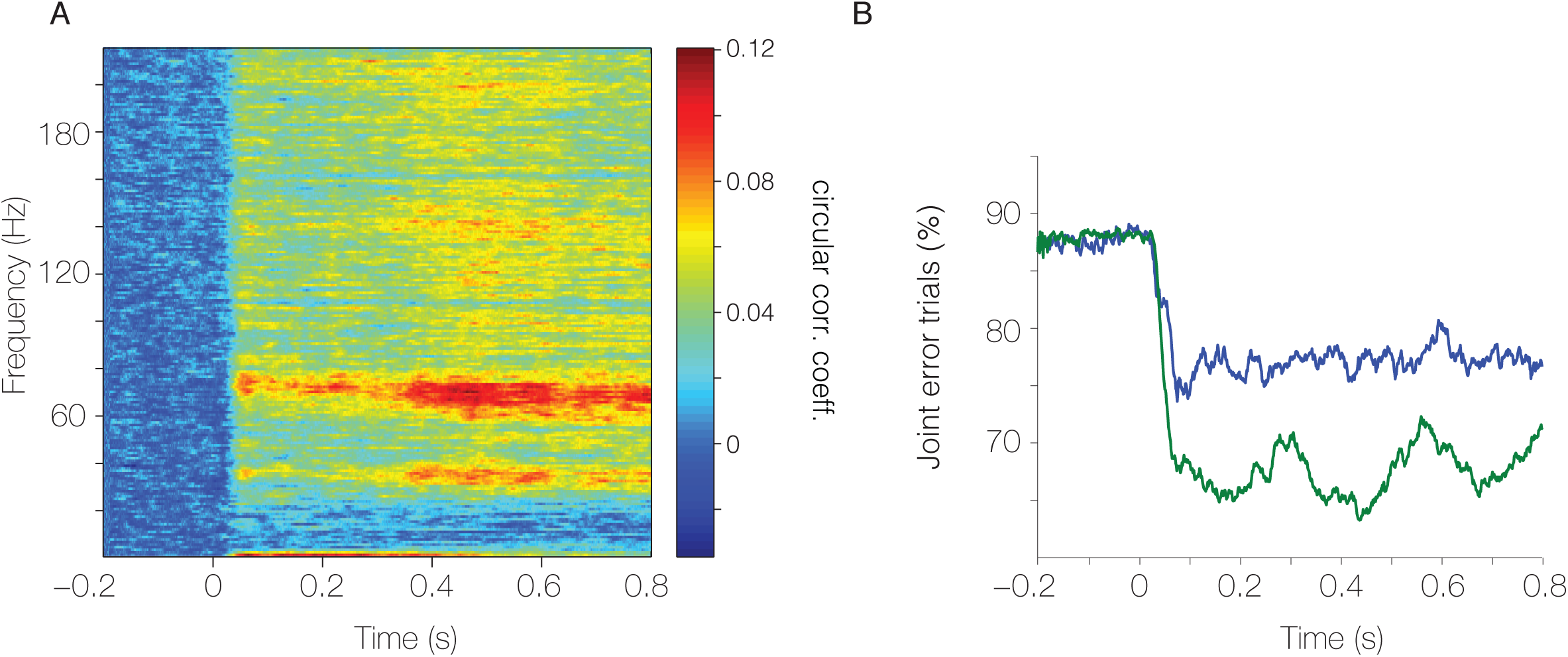
Trial-wise comparison of LFP and MUA decoding performance. (A) Correlation of trial-by-trial estimation of stimulus orientation based on LFP power and MUA rate. Both signals show a modest degree of correlation peaking in the frequency bands of highest accuracy for the LFP power. (B) Joint error trials. The percent of trials in which LFP (green) or MUA (blue) misclassification occurred together with an error in the simultaneously recorded complementary signal. Errors in LFP and MUA often occurred in unison. Results show population values across sessions and monkeys.

### Retinotopic selectivity of ECoG signals

We next evaluated the amount of stimulus position information available from the ECoG recordings. The position of all recording sites from the ECoG arrays of both monkeys are presented as projected on a template brain reconstruction in the F99 atlas space (Gonzalez Andino et al., 2007; Van Essen, 2004). To assess retinotopic selectivity across all sites of the ECoG grid, monkeys kept fixation for several seconds while visual stimuli were presented randomly interleaved at 60 different positions in the lower right visual quadrant (Fig 4B), corresponding to the portion of visual space covered by our ECoG. Stimulus-position dependent changes in spectral power (in any frequency band) were established through an Analysis of Variance (ANOVA). The resulting *p*-values are shown in Fig. 4C (for both monkeys combined) and reveal that selectivity for stimulus position was mainly found in areas V1/V2 and areas V4/TEO (Belitski et al., 2008; Berens et al., 2008; Besserve et al., 2015; Jia et al., 2011; Kayser and König, 2004; Lewis et al., 2016; Myers et al., 2015). We also determined the degree to which recording sites contained information about visual eccentricity (Fig. 4D) or polar angle (Fig. 4E). The sites containing visual information were stable across absolute position, eccentricity and polar angle. We selected those sites for further analyses (V1/V2: 68 sites in monkey P, 32 in monkey K; V4/TEO: 17 sites in each monkey).

**Figure 4.**
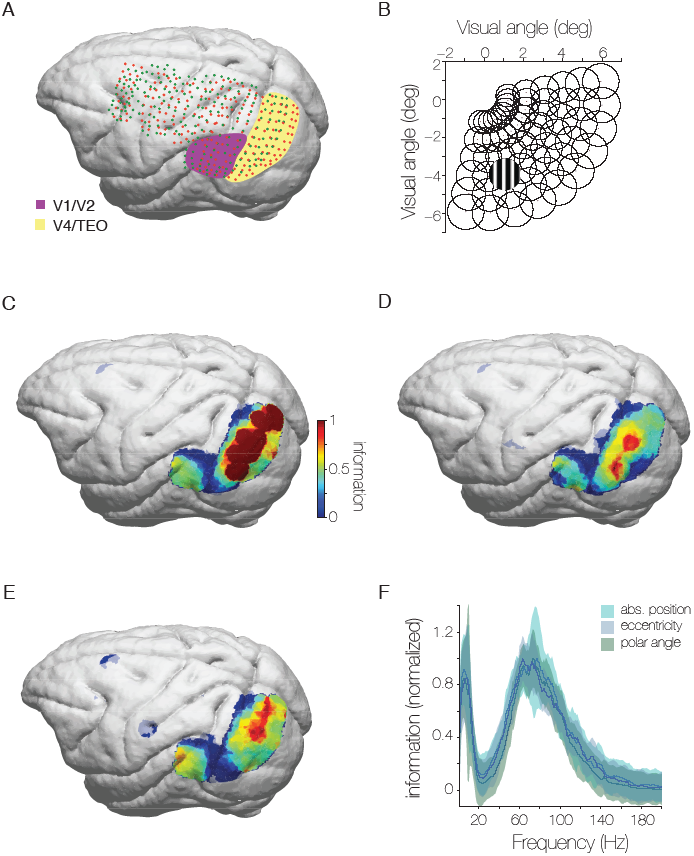
High density ECoG layout, receptive field mapping paradigm and spatial specificity of visual information from LFP. (A) Rendering of the ECoG grids of monkeys K and P overlaid on an atlas brain in F99 space. Lines indicate the covered area with the major sulci. Dots indicate the 218 bipolar electrode derivations. Sites considered as lying in areas V1 and V2 are highlighted in yellow and those in areas V4 and TEO are highlighted in purple. Monkey K in green and Monkey P in red. (B) Receptive fields were mapped with scrolling gratings at 60 locations in the lower right quadrant, corresponding to the coverage of the ECoG array. (C) Selectivity of all ECoG sites for stimulus position based on total stimulus induced power in all frequency bands projected onto atlas brain for monkeys P and K. (D) Same as C, but for eccentricity. (E) Same and C, but for polar angle. (F) Selectivity of visually-selective ECoG sites for absolute position, eccentricity and polar angle as a function of frequency.

### Frequency specificity of positional information

After confirming that power in the LFP could differentiate the position of visual stimuli, we investigated whether this effect was broadband, occurring similarly across many frequencies, or whether it occurred for specific, band-limited rhythms. We limited this analysis to visually driven recordings sites (determined from broadband power). Fig. 4F shows the amount of information about stimulus position decoded from the LFP power as a function of frequency. Different aspects of stimulus position, i.e. absolute position, eccentricity and polar angle, are shown as separate lines. Because decoding of those different aspects was based on different degrees of freedom (60 absolute positions, 6 eccentricity bins, 10 polar angle bins), information is shown after normalizing the peak to unit value. This allows a direct comparison of the spectral shape and demonstrates that different aspects of retinotopy are represented in very similar rhythms, namely in an alpha-beta band and a gamma-frequency band.

### Time-frequency dynamics of positional information and classification error

As decoding of the different aspects of stimulus position led to very similar spectra, we focused on the absolute spatial position as the most general aspect and investigated the stimulus information as a function of frequency and time. As described previously, we applied naïve-Bayes classification to our single-trial power estimates in order to predict stimulus position given the power at a specific point in time and for a specific frequency. Fig. 5A shows the accuracy values as a function of time and frequency, illustrating that e.g. LFP gamma power allowed accurate prediction of the absolute stimulus position in 30–40 % of trials. Chance level is 1/60 (1.67%). Our results were highly reproducible across a broad range of training set sizes. Further, for each time-frequency point, we repeated training and classification 100 times on randomly selected sets of trials in order to insure our classification was robust to variations in the training and test sets. While stimulus position information in the alpha-beta band is primarily transient, there is both a transient and a sustained component in the gamma band. The transient performance was more broadband, while the sustained performance was more band-limited, but maintained stimulus information in a continuous manner. This is further illustrated in the respective time courses shown separately in Fig. 5B. Fig. 5B also shows the same analysis for the broadband LFP amplitude.

**Figure 5.**
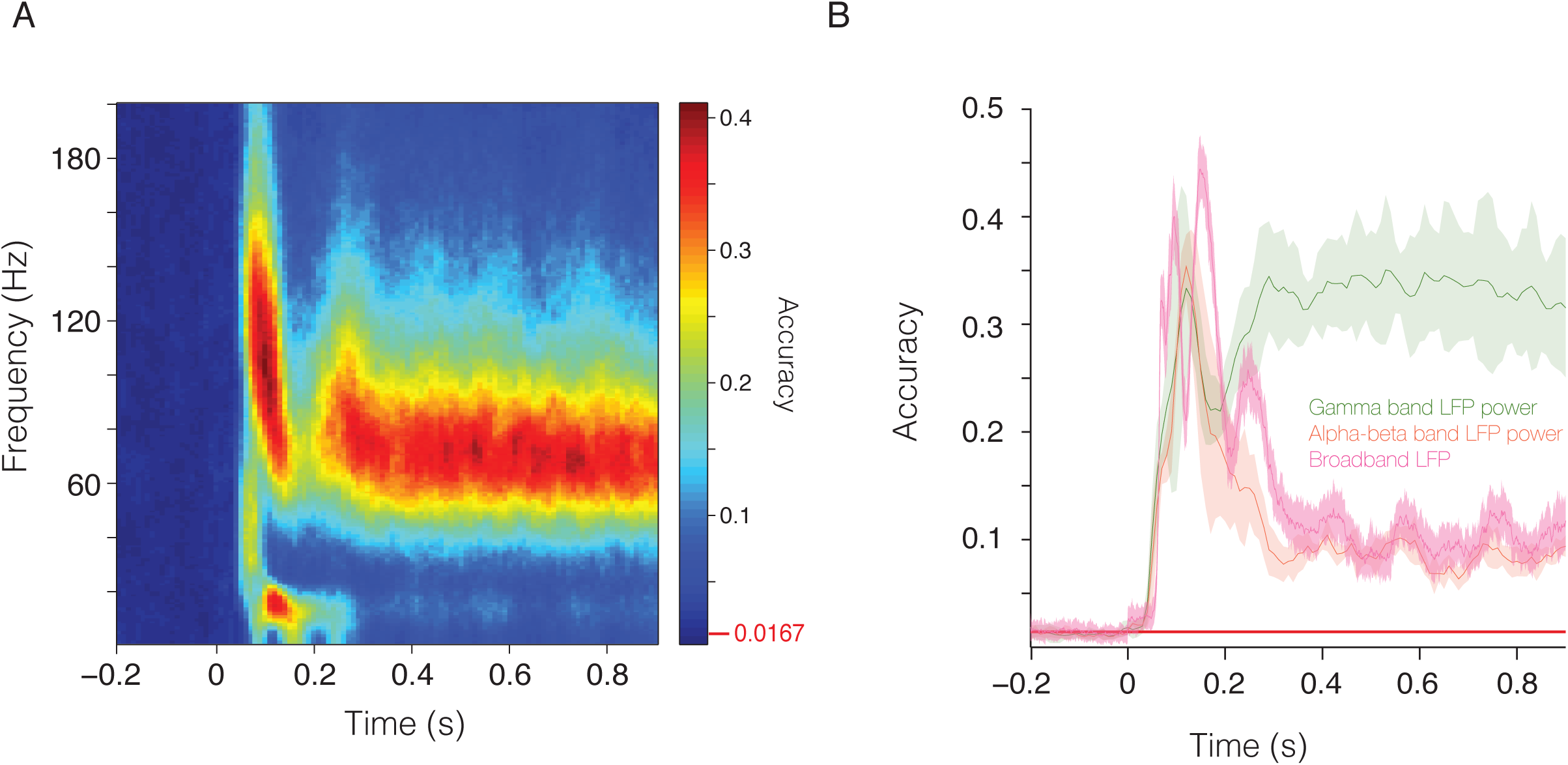
Frequency and temporal specificity of stimulus position information from LFP. (A) Time-frequency plot showing position decoding accuracy for a decoder trained separately on each time-frequency combination. Peak decoding performance is reached by 100 ms. The red line indicates chance level performance. (B) Time course of mean accuracy for gamma band LFP power (green), alpha-beta band LFP power (orange), or broadband voltage (pink) for stimulus position as a function of time. The red line indicates chance level performance. Results are the average performance across randomized train/test sets for monkeys K and P.

We next investigated the size of errors made during misclassifications. This was important because a well-behaved decoder should make minimal errors in classification if the information contained in the feature set is well ordered based on the characteristic of interest. The classification performance for an example stimulus can be seen in Fig. 6A. The veridical stimulus was presented in the location marked by the white asterisk. The circular outlines represent the size of the different decoded stimuli and the red circles highlight the relative proportion the decoder identified the respective location. The fact that misclassifications occur in a clustered fashion around the veridical stimulus location and the fact that they mostly occur for stimuli that overlapped with the veridical stimulus suggest that the position information available in the LFP is highly structured. In order to assess this scatter across all stimulus positions, we calculated the mean overlap between the veridical stimulus position and the position estimated by the decoder. We computed the overlap as the mean of the intersection between the areas of the veridical and error stimuli as a percentage of the veridical stimulus area for each stimulus position. The results of this analysis are presented in Fig. 6B and indicate that errors typically overlapped with the veridical stimulus by an average of 25% in the two frequency bands with highest accuracy. In addition to intersection, we could quantify errors by the distance of the center of the estimated stimulus position from the center of the veridical position. We calculated the distance between the positions for all misclassified trials and found that within the two frequency bands of highest accuracy, the mean distance of error trials was less than 1.5 degrees of visual angle.

**Figure 6.**
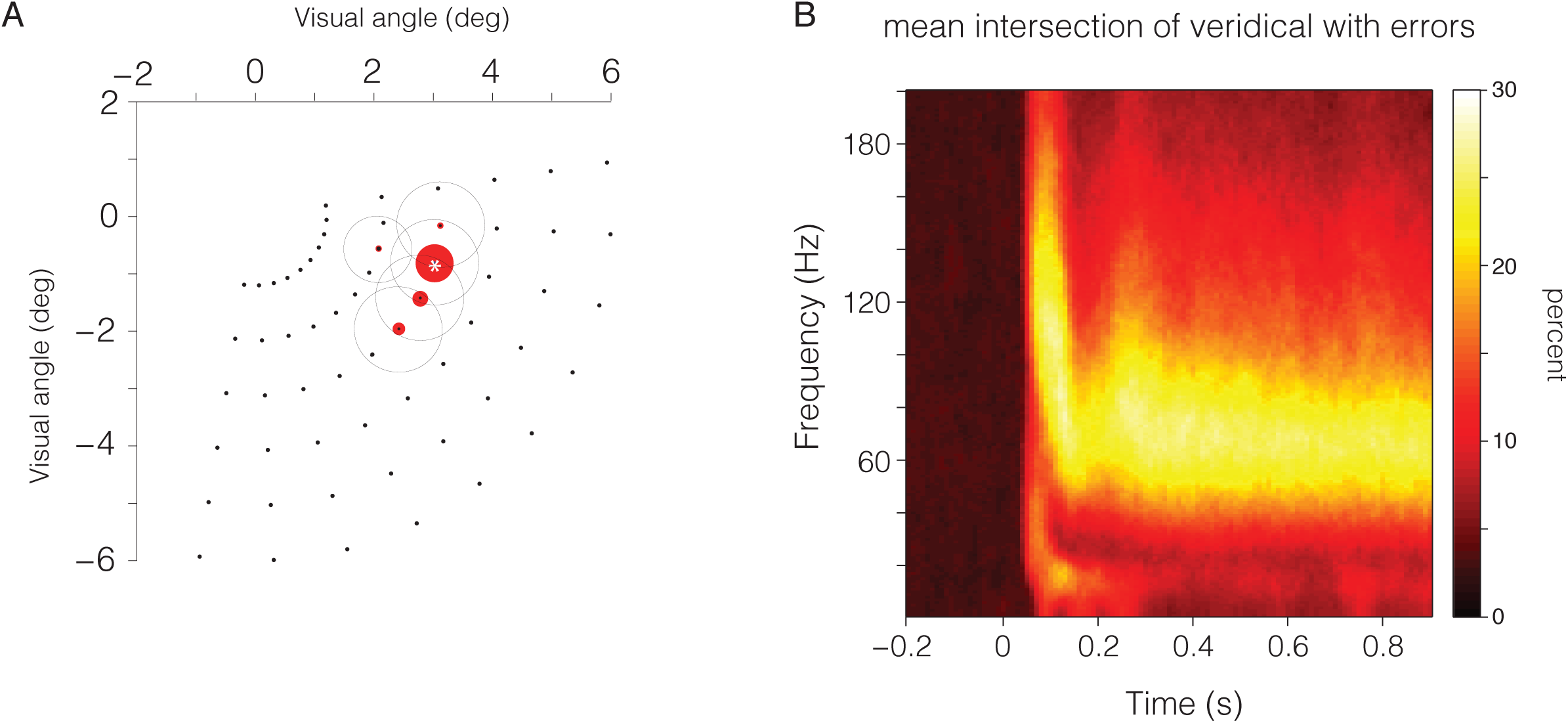
Time-frequency resolved stimulus decoding and errors. (A) Example decoding performance, showing that misclassifications are spatially close and often overlapping with the veridical stimulus. Red dots show the stimulus-position likelihoods estimated based on a stimulus shown at the position marked by the white asterisks. Example is from monkey P. (B) Time-frequency plot showing overlap (in percent) across all misclassified trials of the estimated position and the veridical position across all stimulus positions. Errors in position estimation tended to overlap with the veridical stimulus position demonstrating the high degree of accuracy and spatial clustering of estimates. Results are for monkeys K and P.

### Spatial pattern of positional decoding weights

To further understand the operation of our decoder, we investigated the spatial pattern of the read-out weights across V1 and V4. The decoder uses recording sites as features and fits a probability distribution for each recording site and stimulus position. After decoder training, each stimulus position is associated with a weight matrix assigning each recording site a value, the magnitude of which determines how a specific site contributes to the coding of a specific stimulus position. Decoding of a given stimulus position typically relied on a locus of recording sites reflecting the underlying retinotopic organization. The read-out weights for a specific stimulus location and all time-frequency bands of high accuracy are shown in Fig. 7A. Because the read-out weights of our decoder correspond to the degree to which LFP power from a recording site is able to provide useful information about stimulus position, we reasoned that our decoder could be tapping into the retinotopic organization of visual cortex in estimating stimulus position. Therefore, in order to compute the retinotopy of the ECoG recordings from V1, V2, V4 and TEO, we grouped the 60 stimulus locations according to eccentricity or polar angle. For each recording site, we assigned the eccentricity and polar angle of the stimulus position for which that site had the largest read-out weight. The eccentricities and polar angles of those stimulus positions are shown for each recording site in Fig. 7B and C. Both eccentricity (Fig. 7B) and polar angle (Fig. 7C) were represented in orderly retinotopic maps, that corresponded well with previously determined topographies from repeated recordings with penetrating electrodes (Gattass et al., 2005) or from fMRI (Brewer et al., 2002). Two contiguous maps of space were visible, one behind the lunate sulcus for areas V1/V2, and another one between the lunate and the superior temporal sulcus for areas V4/TEO. For simplicity, we will refer to ECoG sites in the V1/V2 map as V1, and to sites in the V4/TEO map as V4. While such an ordering may seem trivial, the fact that decoding weights reflected areal organization speaks to both the local nature of the recorded signals, as well as the interpretability of our decoding mechanism. While machine-learning algorithms can often lead to results that are hard to interpret, our decoder seems to correspond to the functional organization of the underlying cortical areas.

**Figure 7.**
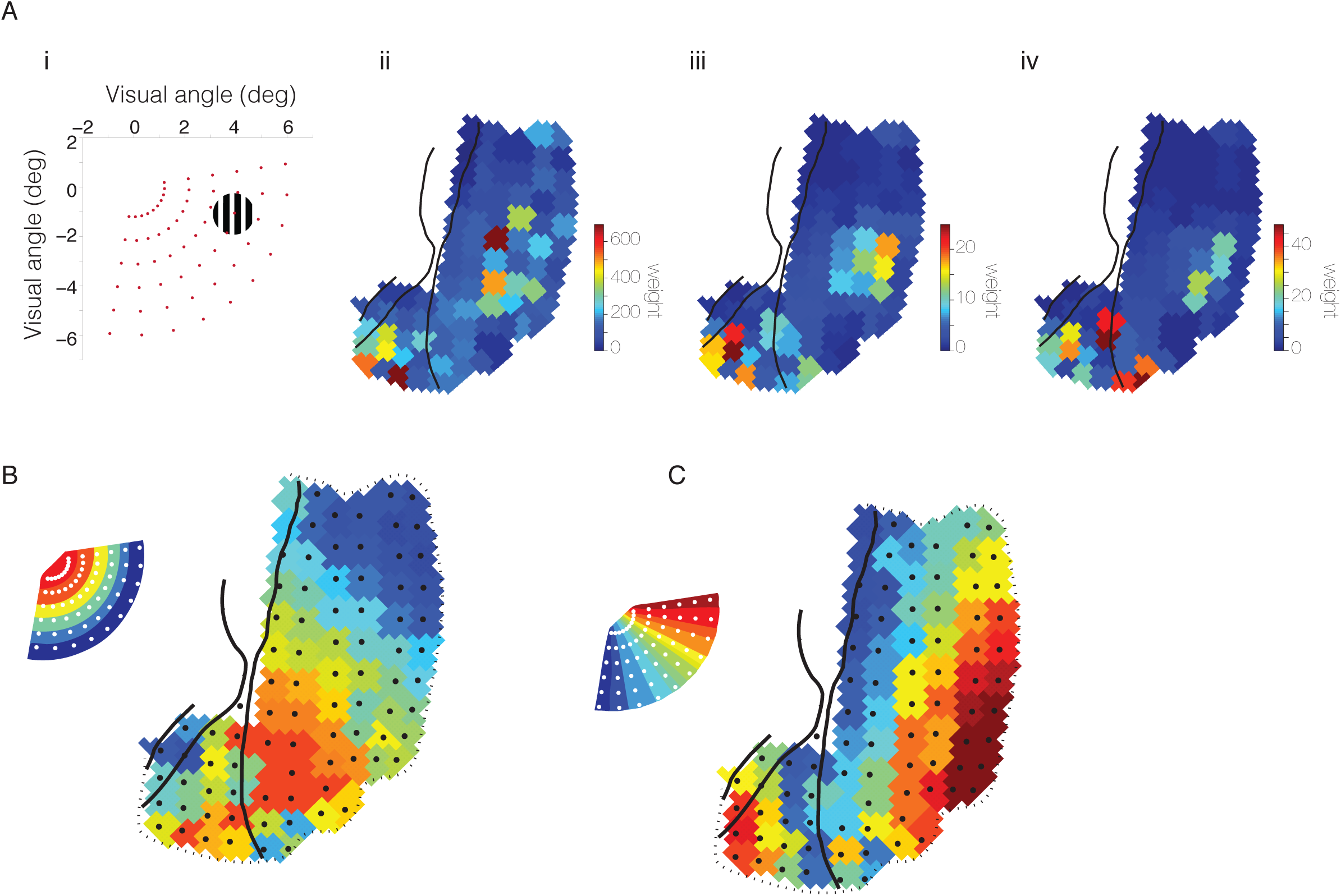
Retinotopic maps based on gamma band (80-95 hz) activity in monkey P. (A) Decoding weights reflect loci of high dependence for a single stimulus position. (i) Example stimulus position. (ii) Map of decoder weights for the low frequency power (80-150ms and 10-20Hz) for the position shown in (i). (iii) Same as in (ii), but for transient gamma band power (80-150ms and 80-130Hz). (iv) Same as in (ii), but for sustained gamma band power (280-900ms and 60-86Hz). (B and C) Retinotopic maps computed from decoding weights across all positions. (B) Map of eccentricity; each recording site is colored to indicate the mean eccentricity of the 5 stimuli giving the largest gamma band response. (C) Map of polar angle; each recording site is colored to indicate the mean polar angle as estimated above. Inset shows how the 60 stimulus locations are represented across eccentricity and polar angle.

### Subsampling performance and the accuracy of single sites

We next sought to understand the scaling of decoding performance with the number of recorded sites. To this end, we investigated classification accuracy as a function of the number of recording sites made available to the decoder. The results of our recording site subsampling are displayed in Fig. 8A for area V1 and in Fig. 8B for area V4. Performance in V1 was significantly higher than that in V4, even when considering a V1 decoder trained on the smaller number of recording sites available for V4 (17 sites, 36.26% in V1 versus 17.55% in V4, p < 10^−8^, two-sided t-test). This indicates that position is more accurately represented by LFP in V1, which is likely a consequence of the difference in receptive field size between the two areas, as well as differences in areal size and magnification factor, leading to differences in spatial resolution.

**Figure 8.**
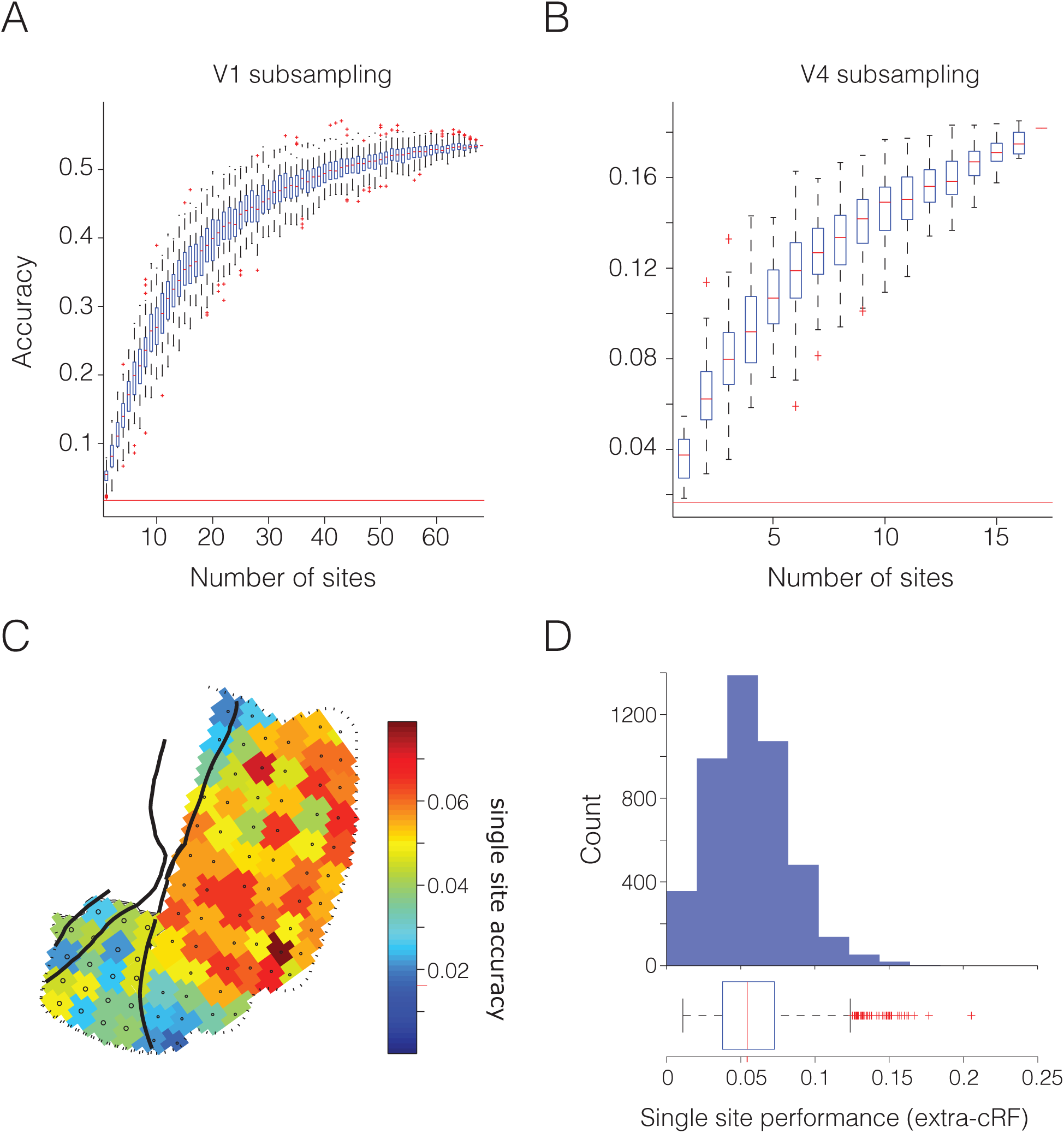
Subsampling performance and single channel decoding. (A) Effect of classifying stimulus position based on subselected channels in V1. (B) Same as A, but for V4. (C) Performance of decoder trained to distinguish position based on single channels. Single V1 channels have higher performance than V4. Single channels contain a significant amount of information about stimuli lying outside of the classical receptive field. (D) Performance of single channels in V1 restricted to the 20 stimuli > 2.5 degrees from the classical receptive field. Mean performance across all channels is higher than chance and many channels show performance 2-3x that expected by chance. Results are from monkeys K and P.

In both areas, we were able to perform significantly above chance level using only the information from a single recording site. The performance of individual recording sites in both areas are shown in Fig. 8C. Again, the increased accuracy of positional information in V1 is evident from the single recording site analysis. These results suggest that V1 neurons, while being retinotopically organized, nevertheless carry information about visual stimulation occurring far from their classical receptive field. Alternatively, high single recording site performance could be due to the subset of stimuli lying near the classical receptive field, i.e. in Fig. 4B, the set of stimuli having a non-zero intersection. In order to decide between these options, we performed an analysis for single recording site performance restricted to stimuli more than 2.5 visual degrees outside of the classical receptive field defined for each site. For this analysis, the number of stimuli was limited to 20 and we used the 45 most or least eccentric V1 recording sites. As shown in Fig. 8D, many single channels were able to determine the position of the stimulus with performance 2-3x greater than expected by chance. The maximally performing channel achieved four times chance performance. This was also true of the average across all tested recording sites (mean single-channel performance different from chance, two-sided t-test, p=0.003). The fact that individual channels have information that allows relatively accurate performance across the sampled visual quadrant suggests that the LFP may reflect more global aspects of stimulation.

### Classification of natural scene and object identity from LFP

Finally, we investigated the presence of stimulus specific patterns of LFP power in V1 and V4 during viewing of natural images. We estimated time-resolved single-trial power for V1 and V4 sites acquired while two monkeys viewed a static natural image. We limited our analysis to images that were repeated at least 10 times so that we had enough stimulus repetitions to train a decoder. The monkeys fixated while waiting for a natural image to appear and we were able to determine the identity of the image with peak accuracy above 60% (chance level = 1/16 (6.25%)). The results of the time-frequency resolved decoding of natural image identity are presented in Fig. 9. Critically, the same frequency bands that had the greatest position, orientation and direction information also had the greatest information about natural scene and object identity. Analysis was limited to the period starting 200 ms prior to stimulus onset and proceeding up to the first saccade. This was necessary due to the trial design: prior to stimulus presentation, monkeys fixated a small dot in the center of the screen, resulting in a stable image position across trials. However, after stimulus onset, they were free to move their eyes, and they typically initiate a saccade away from the fixation point after 200-300 ms. It was not possible to classify images after the initial saccade, because each saccade moved the image on the retina in an unpredictable manner, not reproduced across trials. The fact that similar frequency bands contain a high degree of stimulus specific information also during natural image viewing, suggests that the spatial pattern of band-limited activity across whole cortical areas accurately encodes the identity of visual scenes.

**Figure 9.**
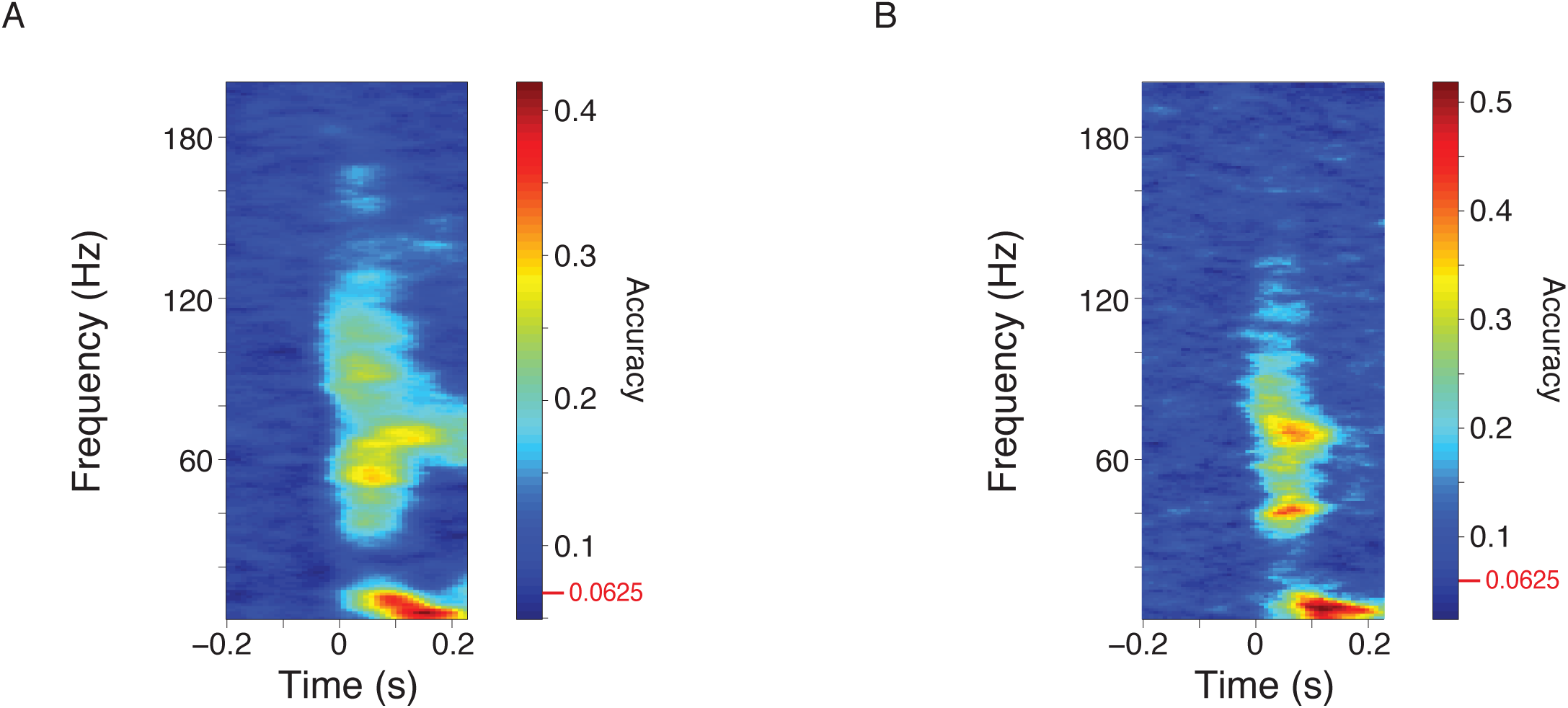
Identification of natural scenes and objects from ECoG. (A) Time-frequency classification performance for color natural images presented to two monkeys. The red line indicates chance level performance. (B) Time-frequency classification performance for black and white natural images presented to two monkeys. The red line indicates chance level performance. Results are the average performance across randomized training/test sets for monkeys P and A.

## Discussion

In summary, we found two frequency bands that reliably contained the most stimulus-related information across a variety of stimulus features, viewing contexts, recording techniques, and monkeys. By combining datasets from intracortical microelectrode recordings and from large-scale electrocorticography, we were able to demonstrate the robust existence of stimulus-specific information in the band-limited power of the LFP. By comparing the performance of decoders trained on the population rate vector and those trained on the band-limited power of LFP, we were able to directly compare stimulus-related information available from the LFP to that in the MUA. Overall, similar amounts of stimulus-related information were in the alpha-beta and gamma bands as was in the spike rate. Further, in the case of stimulus-position decoding, the pattern of activity learned by the trained decoders was organized according to areal topography, suggesting that these weights might provide a means to construct functional maps in early sensory areas and potentially also higher order areas (Mante et al., 2013; Rigotti et al., 2013). The fact that decoding of both, rhythmically synchronized activity, as captured by the LFP power, and decoding of spike rates showed considerable stimulus selective information during the initial transient indicates that higher-level areas, which may use these signals in order to extract incoming information, can differentiate the parameters of stimulation on behaviorally relevant time scales (VanRullen et al., 2005). Finally, the fact that we could distinguish image identity from band-limited LFP during viewing of natural images suggests that the spatial distribution of LFP power across cortical areas may reflect the coordination of cortical activity into a cohesive representation of the sensory environment. We found the greatest amounts of stimulus-related information in two distinct frequency bands across multiple low-level visual features, as well as natural image identity. These two bands, the alpha-beta and the gamma band, most likely reflect different sources. Due to the temporal profile, as well as the frequency content of the alpha-beta band, it likely reflects the visual evoked potential (Schroeder et al., 1995). Gamma band power showed two distinct patterns of information-rich activity across stimulus features, recording techniques and viewing conditions. One was relatively broadband (~ 60-160Hz), with short latency, and a brief duration. This broadband gamma was followed by a gamma that was narrower-band, had a longer latency and was sustained throughout stimulus presentation for most features and contexts. The transient, broadband gamma activity likely reflects a mixture of rhythmic and non-rhythmic components. The rhythmic component reflects the same gamma rhythm that is also present during the sustained phase, but that is of higher frequency after the salient stimulus onset (Fries, 2015). This gamma rhythm might vary in breadth on every trial, as bandwidth often scales with peak frequency (Buzsáki, 2006). In addition, its peak frequency might change with changes in stimulus and the animal’s state (Jia et al., 2013). Non-rhythmic components that might contribute are postsynaptic potentials (PSPs) impinging onto the recorded neurons, and action potentials (APs) generated by those neurons. PSPs show an exponential decay, corresponding to a 1/f spectral signature (Einevoll et al., 2013; Miller et al., 2009). Extracellularly recorded APs extend over 1-3 ms, corresponding to a spectral power peak around 1-2 kHz, yet extending down into the LFP frequency range (Ray et al., 2008). We estimate that the genuinely non-rhythmic components play a minor role in our LFPs from microelectrodes and ECoG, because stimulus decoding was most successful for the spectrally limited gamma band.

The decoding of natural stimulus identity succeeded for power in a short time period and a relatively broad spectral band. This broad spectral band likely does not reflect an early, transient peak of neuronal PSPs and APs. PSPs lead to 1/f power and therefore, decoding carried by PSPs should also exhibit a 1/f spectral distribution, which is not consistent with Fig. 9. Similarly, APs lead to power that increases up to approximately 1-2 kHz, and decoding based on APs should correspondingly show this spectral shape, again inconsistent with what we found. The smooth drop-off in decoding performance from the gamma band towards 200 Hz is not due to filtering, because the hardware filter was at 8 kHz and the offline LFP filter was at 250 Hz and relatively sharp. Rather, we suggest that decoding of natural stimulus identity has a relatively broad spectral characteristic, because natural stimuli show substantial intra-and inter-stimulus variance in stimulus features and probably also in consequent attention, which both affect gamma frequency (Bosman et al., 2012; Fries, 2015; Jia et al., 2013). The relatively short duration of successful image decoding is due to the fact that the animals were fixating only until the stimulus appeared and were free to explore the image with saccades thereafter. In this case, first saccades typically occur around 250 ms and move the image across the retina in different ways in each trial.

Several previous studies have established that stimulus selectivity differs across frequency bands of the LFP power spectrum. Gamma-band power is selective for stimulus orientation (Frien et al., 2000; Kayser and König, 2004; Siegel and König, 2003), and for spatial and temporal frequency (Kayser and König, 2004). Correspondingly, we report that decoding of gamma power from multiple simultaneous recordings allows orientation decoding, and we demonstrate that accuracy is comparable to firing rate decoding and the respective errors are correlated across trials. These decoding results might have been partly predicted by previous studies, yet are also partly surprising. Orientation preferences derived from LFP gamma power and spike rates of the same recording site have been reported to differ from each other, when large stimuli are used (Berens et al., 2008; Jia et al., 2011), with the gamma-power preferences typically being essentially identical for multiple nearby-recorded sites (Berens et al., 2008). Notably, even though the stimuli used here were of similar size, gamma power allowed reliable orientation decoding. Also the decoding of stimulus motion direction from LFP power was not expected. A previous study investigated orientation and direction tuning of LFP and MUA power in different frequency bands (Frien et al., 2000). For orientation, the tuning was significant for both LFP and MUA and particularly sharp for the gamma band. In contrast, none of the recording sites was direction selective for any one of the frequency bands, even though MUA was analyzed, which has a particularly high spatial specificity. When we decode LFP power from three to eight electrodes, accuracy for stimulus direction is well above chance level (2.5x for the gamma band and 3.2x for MUA). Thus, the use of several simultaneously recorded LFPs in conjunction with decoding algorithms seems to provide a more sensitive approach to quantify the information present in measures of neural activity as compared to tuning-curve based approaches. In addition, the spatial specificity of the recorded signal appears to be less relevant than one might assume. The ECoG electrodes used here were 1 mm in diameter and placed merely onto the cortical surface. This is a great advantage for potential BMI applications in which brain damage needs to be minimized and long-term viability is essential. Previous studies demonstrated that ECoG-LFP power allows decoding of stimulus shape (Rotermund et al., 2009) and of the location of selective attention (Rotermund et al., 2013). We find that decoding of ECoG LFP allows a precise localization of small stimuli, a capacity highly relevant for BMI. The frequency ranges carrying information about stimulus eccentricity and polar angle were essentially identical. Intriguingly, loss of some ECoG electrodes did not preclude accurate stimulus localization, as shown by our analysis of recording site subsampling. Recent studies also reported stimulus dependence of phase relations (Agarwal et al., 2014; Lubenov and Siapas, 2009; Maris et al., 2013; Rotermund et al., 2013; van Ede et al., 2015). Yet, decoding of selective attention based on gamma-band phase relations did not exceed decoding based on gamma-band power (Rotermund et al., 2013). Therefore, we focused here on spectral power.

Thus, convergent evidence shows that aspects of rhythmic neuronal synchronization encode stimulus information. Optimal stimulus encoding has been conceptualized within sparse coding theory (Olshausen and Field, 2004; 1996). Sparse coding models place an explicit constraint on the joint activity of sensory cells, without specifying the physiological basis of this constraint. Such a constraint is likely to involve recurrent and specifically inhibitory activity, which in turn constitutes the mechanism behind gamma-band synchronization (Buzsáki and Wang, 2012; Knoblich et al., 2010). It is appealing to consider that oscillatory synchrony, mediated by recurrent inhibitory connections, constrains population activity based on spatial and temporal context through the dynamic balance of excitation and inhibition (Haider et al., 2010; Vinck and Bosman, 2016). In such a scenario, single cells, rather than reflecting the disjoint feature extractors of classical serial feature extraction models, would be conceived of as integral members of the local population (Graf et al., 2011), embedded within the coordinated orchestration of temporal and spatial context enforced by the distributed oscillatory regime (Lewis et al., 2015). In this scheme, aspects of stimulus representation that are classically attributed to firing rate responses in higher visual areas, like complex selectivity in concert with invariance, are already present in the pattern of synchronization in lower visual areas and therefore can be decoded from simultaneous multisite recordings (Singer, 2013). Machine-learning techniques provide a means to explore which information is encoded in cortical networks and how (Kriegeskorte and Kievit, 2013). By iteratively applying the logic of machine learning to multiple areas in sensory hierarchies, we can begin to tease out how information is represented and transformed in the brain. Such approaches have already begun to reveal the formation of invariance along ventral stream visual areas (DiCarlo et al., 2012; Freiwald et al., 2009; Hong et al., 2016; Li and DiCarlo, 2012; Pagan et al., 2013). Likewise, when applied to higher order areas, machine-learning methods may help us develop maps and canonical computations for previously intractable association areas. More work is needed to determine the extent to which the spatial distribution of LFP reflects bottom-up features of the sensory stimulus (Lowet et al., 2015), such as local indices of contrast, orientation, or luminance, and to which extent it reflects lateral or feed-back activity which constrains the bottom up signal into a fixed percept (Bastos et al., 2015). It will also be important to determine the extent to which the information contained in spike rates and LFP are redundant and the degree to which they carry synergistic information, though some work already suggest they are non-redundant (Belitski et al., 2008) and possibly synergistic (Womelsdorf et al., 2012). Using measures of spiking separate from rate could also add substantial information. The specific sequence of spikes across the population, spike-LFP phase (Vinck et al., 2010), or even the distribution of the inter-spike-interval for single units may provide additional information about stimulus parameters, which may impact downstream areas. Future studies with simultaneous measurement of activity within local and distributed populations, especially those with coverage at multiple spatial scales and positions in the sensory hierarchy (Lewis et al., 2015), will increase our understanding of how populations of cells interact to flexibly represent a dynamic world.

## Materials and Methods

### Intracortical microelectrode recordings

#### Techniques and Signal Preprocessing

All procedures for the 3 monkeys (J, L and N) with quartz-insulated platinum/tungsten microelectrode recordings were approved by the German local authorities (Regierungspräsidium, Hessen, Darmstadt), were in full compliance with the guidelines of the European Community for the care and use of laboratory animals (European Union directive 86/ 609/EEC) and have been previously published (Lima et al., 2010; Vinck et al., 2010; Womelsdorf et al., 2012). Experiments were performed on three awake adult monkeys as previously described in detail (Lima et al., 2010; Vinck et al., 2010; Womelsdorf et al., 2012). Recordings were performed in area V1 using 1-10 tungsten/platinum electrodes. Monkeys passively viewed drifting gratings of varying orientations (16 directions in steps of 22.5 deg). Before the experiment, each monkey was surgically implanted with a head post, a scleral search coil, and a recording chamber. Recordings were made from the opercular region of V1 (receptive field centers: 2-5.8 deg of visual angle in eccentricity) and from the superior bank of the calcarine sulcus (receptive field centers: 8-12.8 deg of visual angle in eccentricity). Recordings proceeded with electrodes inserted independently into the cortex through transdural guide tubes with precision hydraulic micro-drives mounted onto an X-Y stage (MO-95; Narishige Scientific Instrument Laboratory). Spiking activity and the LFP were obtained by amplifying (1,000×) and band-pass filtering (multiunit activity: 700-6,000 Hz; LFP: 0.7-170 Hz) the recorded signals using a customized 32-channel headstage and preamplifier (headstage HST16o25; headstage and preamplifier from Plexon Inc.). Additional 10× signal amplification was performed by onboard amplifiers (E-series acquisition boards; National Instruments). LFPs were acquired with a resolution of 1.0 ms. Spikes were detected online by an amplitude threshold. Spike events and corresponding waveforms were sampled at 32 kHz, and spike waveforms were recorded for 1.2 ms. The MUA used in this study consisted of all of the binarized, threshold-crossings on a given electrode, after removing artifacts.

#### Visual Stimulation and Behavioral Task

Stimuli were presented as movies at 100 or 120 frames per second using a standard graphical board (GeForce 6600 series; NVIDIA). The CRT monitor used for presentation (CM813ET; Hitachi) was gamma corrected to produce a linear relationship between output luminance and gray values, and subtended a visual angle of 36 × 28.8 deg (1,024 × 768 pixels). At the beginning of each recording session, receptive fields were mapped using an automatic procedure, in which a bar was moved across the screen in 16 different directions (160 trials) (Fiorani et al., 2014). Receptive field position was estimated from the global maximum of a response matrix, at a resolution of ~6 min of arc. Subsequently, monkeys passively viewed drifting gratings during fixation of a small central fixation spot. Gratings had spatial frequencies ranging from 0.5 to 2.0 cycles per degree and velocities ranging from 0.5 to 3.0 degrees per second. Grating drift directions were generated randomly from a total of 16 directions (steps of 22.5). The stimuli were centered over the receptive fields within a circular aperture of 8.08 degree. After the monkey acquired fixation, there was a pre-stimulus baseline of 800-1,000 ms, after which the stimulus was presented for 800-1,400 ms. To obtain a reward, monkeys had to release the lever within 500 ms after the fixation point had changed color. Trials were aborted upon fixation breaks, or when the lever was released before the color change. Eye position was monitored continuously by a search coil system (DNI; Crist Instruments) with a temporal resolution of 2 ms.

### ECoG recordings

#### Techniques and Signal Preprocessing

All procedures for the 3 monkeys (K, P and A) implanted with ECoG arrays were approved by the ethics committee of the Radboud University, Nijmegen, NL and have been previously published. Neuronal recordings were made from three left hemispheres in three monkeys through a micro-machined 252-channel ECoG array implanted subdurally (Bosman et al., 2012; Brunet et al., 2013; Rubehn et al., 2009). Briefly, a ~6.5 x ~3.5 cm craniotomy over the left hemisphere in each monkey was performed under aseptic conditions with isoflurane/fentanyl anesthesia. The dura was opened and the ECoG was placed directly onto the brain under visual control. Several high-resolution photos were taken before and after placement of the ECoG for later co-registration of ECoG signals with brain regions. After ECoG implantation, both the bone and the dural flap were placed back and secured in place. ECoG electrodes covered numerous brain areas, including superficial parts of areas V1, V2, V4 and TEO. As mentioned in the main text, retinotopic mapping revealed two contiguous maps of space, one behind the lunate sulcus for areas V1/V2, and another one between the lunate and the superior temporal sulcus for areas V4/TEO. For simplicity, we refer to ECoG sites in the V1/V2 map as V1, and to sites in the V4/TEO map as V4. After a recovery period of approximately 3 weeks, neuronal recordings commenced. Signals obtained from the electrode grid were amplified 20 times by eight Plexon headstage amplifiers, then low-pass filtered at 8 kHz and digitized at 32 kHz by a Neuralynx Digital Lynx system. LFP signals were obtained by low-pass filtering at 250 Hz and down-sampling to 1 kHz. Power-line artifacts were removed by digital notch filtering. The actual spectral data analysis included spectral smoothing that rendered the original notch invisible.

#### Visual Stimulation for Receptive Field Mapping

Stimuli and behavior were controlled by the software CORTEX. Stimuli were presented on a cathode ray tube (CRT) monitor at 120 Hz non-interlaced. When the monkey touched a bar, a gray fixation point appeared at the center of the screen. When the monkey brought its gaze into a fixation window around the fixation point (0.85 degree radius in monkey K; 1 degree radius in monkey P), a pre-stimulus baseline of 0.8 s started. If the monkey’s gaze left the fixation window at any time, the trial was terminated. Several sessions (either separate or after attention-task sessions) were devoted to the mapping of receptive fields, using 60 patches of drifting grating, as illustrated in Fig. 4B. Gratings were circular black and white sine waves, with a spatial frequency of 3 cycles/degree and a speed of 0.4 degrees/s. Stimulus diameter was scaled between 1.2 and 1.86 degrees to partially account for the cortical magnification factor. Receptive field positions were stable across recording sessions (Bosman et al., 2012).

#### Visual Stimulation for Natural Images

In separate recordings sessions, two of the animals (monkey P and A) were required to fixate for 0.63 s on a fixation point (0.12 by 0.12° black square) centered on a gray background, after which a natural image was presented, which was again centered on the background screen. We used 49 grayscale images and 16 color images, with grayscale and color images presented in separate sessions. Grayscale images subtended 16-by-16°, and color images 18.5-by-18.5°. Grayscale images were shown for 3.5-6 s (flat random distribution), and color images for 1.5 s. Once the image had appeared on the screen, the monkey could view it freely. If the monkey kept its gaze on the image as long as it was presented, it was given a juice reward after stimulus offset. Because this task was very easy for the monkeys, almost every trial was rewarded. Each grayscale image was presented for an average of 15 trials, and each color image for an average of 22 trials. Eye position was recorded with an infrared camera system (Thomas Recording ET-49B system) at a sampling rate of 230 Hz. Unless stated otherwise, we selected electrodes over V1 and V4 that were strongly driven by stimuli within the central 4° of eccentricity. Due to placement of the ECoG grid onto the dorsal parts of V1 and V4 in the left hemisphere, receptive fields were in the lower right visual quadrant. Correspondingly, we accepted analysis epochs when the gaze of the monkey was at least 4° away from the lower and the right border of the natural image for at least 90% of the epoch duration. This ensured that the responses of the recorded sites were due to the natural image rather than the screen background or the edge of the image. In monkey P (monkey A), we used 43 (42) electrodes on V1 and 16 (14) on V4. The assignment of electrodes to visual areas was based on intraoperative photographs and brain atlases, and used primarily sulcal landmarks. For most of the electrodes, the area assignment was unequivocal. Yet, some of the most anterior electrodes assigned to V1 might as well be over V2, and the most lateral electrodes assigned to V4 might as well be over the temporal-occipital area (TEO). Exclusion of those electrodes left the results qualitatively unchanged.

### Data Analysis General

All analyses were performed in MATLAB (MathWorks) using FieldTrip (Oostenveld et al., 2011) (http://fieldtrip.fcdonders.nl). For the ECoG datasets, we calculated local bipolar derivatives, i.e., differences (sample-by-sample in the time domain) between LFPs from immediately neighboring electrodes. We refer to the bipolar derivatives as “sites.” Bipolar derivation removes the common recording reference, which is important when analyzing power correlations and/or coherence. Subsequently, per site and individual epoch, the mean was subtracted, and then, per site and session, the signal was normalized by its standard deviation. These normalized signals were pooled across sessions with identical stimulus and task, unless indicated otherwise. In order to select visually selective recording sites, an ANOVA was computed across frequencies. Time-frequency resolved power was computed for the LFP by multiplying the time-domain data by a Hann taper in over-lapping windows of a length scaled to the frequency of interest (3 cycles/window) in step sizes of 1 Hz and 10 ms.

### Spike Rate

All MUA analysis was performed on the spike density calculated from all threshold-crossings on a given electrode. Spike density was computed on a single-trial basis by convolving the binary spike time-series by a Gaussian kernel with a standard deviation of 10 ms. Each trial consisted of 3-9 individual time-series from independent electrodes. Each binary time-series was separately convolved with the kernel in order to derive a continuous variable, which divided by the time window, led to a time-varying spike density estimate.

### Naïve-Bayes Decoding

Single-trial estimates of power or spike density for each condition were used to train a probabilistic decoder using the Gaussian naïve Bayes algorithm. Naïve Bayes applies Bayes’ Theorem with the assumption that features are independent in order to estimate P(X|Y), the probability of feature X, given condition Y. The feature vector was composed of one time window (for MUA decoding) or time-frequency (for LFP decoding) across recording sites. For the Gaussian case, the decoder estimates a mean and standard deviation of each feature (recording site) and each condition (orientation, direction, position, or image identity). When the trained distributions are applied to previously unseen data, the decoder calculates the likelihood associated with each of the possible feature classes (8 orientations, 16 directions of motion, 60 stimulus positions, or 16 natural images). Based on the distribution of likelihoods, we chose the maximum value and used it to quantify classification accuracy. For a given spatial pattern of MUA, LFP power, or broadband voltage, at a specific time, this resulted in a single selection of the most likely class for the given feature. This results in each trial being assigned an expected condition under the learned Posterior, as well as a vector of log-likelihood values for each possible class. For the orientation, direction of motion, and position decoding analysis, our training set consisted of 75% of the available trials for the condition with the fewest number of trials and quantified the performance on the remaining 25% of trials. This avoided introducing *prior* biases into our decoder. We trained on 50% and 90% of the available data and this did not qualitatively change our results. We used an iterated cross-validation approach, which allowed us to control the generalization and robustness of our decoder. Values displayed for the decoding plots shown here display the mean value from 100 iterations of the decoder to avoid any extraneous effects of the training and test sets.

## ACKNOWLEGMENTS

We thank Robert Oostenveld, Eric Lowet, Michaela Klinkmann and Johanna Klon-Lipok for technical assistance during the course of these experiments. This work was supported by DFG (SPP 1665, FOR 1847, FR2557/5-1-CORNET), EU (HEALTH-F2-2008-200728-BrainSynch, FP7-604102-HBP, FP7-600730-Magnetrodes), a European Young Investigator Award, NIH (1U54MH091657-WU-Minn-Consortium-HCP), LOEWE (NeFF).

